# Attention cueing in rivalry: insights from pupillometry

**DOI:** 10.1101/2021.06.27.450072

**Authors:** Miriam Acquafredda, Paola Binda, Claudia Lunghi

## Abstract

We used pupillometry to evaluate the effects of attention cueing on perceptual bi-stability, as reported by adult human observers. Perceptual alternations and pupil diameter were measured during two forms of rivalry, generated by presenting a white and a black disk to the two eyes (binocular rivalry) or splitting the disks between eyes (interocular grouping rivalry). In line with previous studies, we found that subtle pupil size oscillations (about 0.05 mm) tracked alternations between exclusive dominance phases of the black or white disk. These oscillations were larger for perceptually stronger stimuli: presented to the dominant eye or with physically higher luminance contrast. However, cueing of endogenous attention to one of the rivaling percepts did not affect pupil oscillations during exclusive dominance phases. This was in spite of the reliable effects of endogenous attention on perceptual dominance, which shifted in favor of the cued percept by about 10%. The results were comparable for binocular and interocular grouping rivalry. Cueing only had a marginal modulatory effect on pupil size during mixed percepts in binocular rivalry. This may suggest that, rather than acting by modulating perceptual strength, endogenous attention primarily acts during periods of unresolved competition, which is compatible with attention being automatically directed to the rivaling stimuli during periods of exclusive dominance and thereby sustaining perceptual alternations.

**Significance Statement:** Binocular rivalry depends on attention. When it is diverted away from the stimuli, perceptual alternations slow down; when it is preferentially directed to one stimulus, perception lingers more on it, consistent with attention enhancing the effective strength of the rivaling stimuli. Here we introduce pupillometry as a means to indirectly track changes in effective stimulus strength. We find that pupil size oscillates, accurately tracking perceived luminance during two forms of rivalry: binocular rivalry and interocular grouping rivalry. Both show robust effects of attention cueing on perceptual dominance, but pupil oscillations during exclusive dominance are unaffected by cueing. This suggests that endogenous attention does not affect perceptual strength during exclusive dominance, though it might do so during transition phases.

## Introduction

When stimuli in the two eyes are incompatible, binocular fusion fails and perception alternates between the monocular images (binocular rivalry, Wheatstone, 1838). Rivalry has been shown to depend on attention, as perceptual alternations tend to cease when attention is diverted away. When this happens, neural oscillations in early visual areas are also suppressed (Zhang et al., 2011), consistent with the notion that attention modulates the strength of early neural representations (Carrasco, 2011). These effects are adequately modelled by assuming that attentional resources are automatically driven to the dominant stimulus, unless engaged elsewhere; and that attention provides recurrent excitation of the corresponding monocular input, acting synergically with interocular inhibition to maintain the competition between eyes (Li et al., 2017). Besides diverting attention away from the stimuli, cueing attention to one of the rivaling stimuli can affect binocular rivalry, shifting perceptual dominance in favor of the cued percept (Meng and Tong, 2004; Mitchell et al., 2004; Chong et al., 2005; van Ee et al., 2006; Hancock and Andrews, 2007; Paffen and Alais, 2011; Dieter et al., 2016). However, the neural underpinnings of cueing effects have been less systematically studied and, to the best of our knowledge, no previous study has tested whether attention cueing affects the strength of early visual representations during rivalry.

Interocular competition can sometimes be overcome by pattern-based competition, as in interocular grouping rivalry (Alais et al., 2000), where monocular stimuli are complementary, e.g. two half gratings, and perception alternates between images grouped across eyes (Alais et al., 2000). The role of attention in interocular grouping rivalry has not been investigated. In general, attention cueing has more pronounced effects on more complex types of bistable perception, such as Necker cube or bistable structure from motion (Meng and Tong, 2004; van Ee et al., 2005), which could predict stronger attentional modulations in interocular grouping than in binocular rivalry.

Here we propose pupillometry as a method to indirectly index the strength of competing visual representations and objectively quantify the effects of attention cueing on binocular and interocular grouping rivalry.

Pupil size is mainly set by retinal illumination through a simple subcortical circuit (Loewenfeld, 1993). However, light responses are modulated by attention, brightness illusions and contextual processing (Laeng et al., 2012; Binda and Murray, 2015; Mathot, 2018) indicating that the subcortical circuit is fed with cortical signals (Binda and Gamlin, 2017) that represent effective stimulus strength. As long as stimuli are tagged with different luminance, pupil diameter can be used to accurately and precisely track attention in space (Binda et al., 2013; Mathot et al., 2013; Naber et al., 2013) and perceptual alternations over time (Lowe and Ogle, 1966; Einhauser et al., 2008; Fahle et al., 2011; Naber et al., 2011; Turi et al., 2018; Tortelli et al., 2021). Here we exploited this strategy and used luminance to tag pupil responses to stimuli rivaling in perception. We predicted that, if attention cueing enhances the effective strength of the dominant stimulus, pupil oscillations should be amplified. The amplification would provide an objective and time-resolved index of how attention affects binocular and interocular grouping rivalry.

## Materials and methods

### Subjects

We recruited 38 participants (17 males and 21 females including 2 authors, mean age 26.5 ± 0.69 years). Sample size was based on a power analysis that determined the minimum number of participants required to detect a medium sized effect (effect size 0.50, two tailed α 0.05, power 0.8 = 33 participants; we recruited a few more anticipating data losses that fortunately did not occur). Ten additional subjects were recruited for the control experiment (9 females and 1 male, mean age 27.1 ± 0.82 years).

All participants had normal or corrected-to-normal visual acuity (ETDRS charts), normal stereopsis (TNO test) and normal color vision (Ishihara plates); balanced ocular dominance (excluding participants with ocular dominance higher than 70%); no self-reported history of eye surgery, other active eye disease or mental illnesses.

### Ethics Statement

The experimental protocol was approved by the local ethics committee (Comité d’éthique de la Recherche de l’Université Paris Descartes, CER-PD:2019-16-LUNGHI) and was performed in accordance with the Declaration of Helsinki (DoH-Oct2008). All participants gave written informed consent and were reimbursed for the time spent at a rate of 10€ per hour.

### Apparatus, stimuli and procedures

Experiments took place in a dark and quiet room. Visual stimuli were developed in Matlab (The MathWorks Inc., Natick, MA) using Psychtoolbox-3 (Brainard, 1997)running on a PC (Alienware Aurora R8) and a NVIDIA graphics card (GeForce RTX2080). Visual stimuli were displayed on a linearized monitor 53.5 cm-wide, driven at a resolution of 1920 × 1080 pixels. The display was seen through a four-mirror stereoscope which enabled dichoptic viewing of two display areas of 12 × 8 deg each; a chin rest was used to stabilize head position at 57 cm from the display. In each display area, a central red fixation point (0.15 deg in diameter) surrounded by a square frame (3.5 × 3.5 deg) were shown against a uniform gray background (luminance: 12.3 cd/m^2^). The mirrors were carefully adjusted at the beginning of each session to ensure accurate alignment of the dichoptically presented squares. Participants were asked to keep their gaze on the fixation point shown at screen center and to refrain from blinking while the stimuli were on.

Dichoptic presentations consisted of two sets of stimuli, designed to elicit two forms of rivalry: binocular rivalry and interocular grouping rivalry.

For binocular rivalry, visual stimuli consisted of two disks (Figure 1A), 3° in diameter, one white (maximum screen luminance 28.9 cd/m2, Michaelson contrast: 0.31) and one black (minimum screen luminance 1.7 cd/m2, Michaelson contrast: 0.8). Perception alternated between exclusive dominance of the white and the black disk, or mixed percepts (either piecemeal or fusion). In order to discourage fusion, the disks were overlaid with thin orthogonal gray lines (45° clockwise or counterclockwise, 1 pixel wide, corresponding to 0.033 deg, and 0.5 deg apart, with the same luminance as the background).

**Figure 1.**
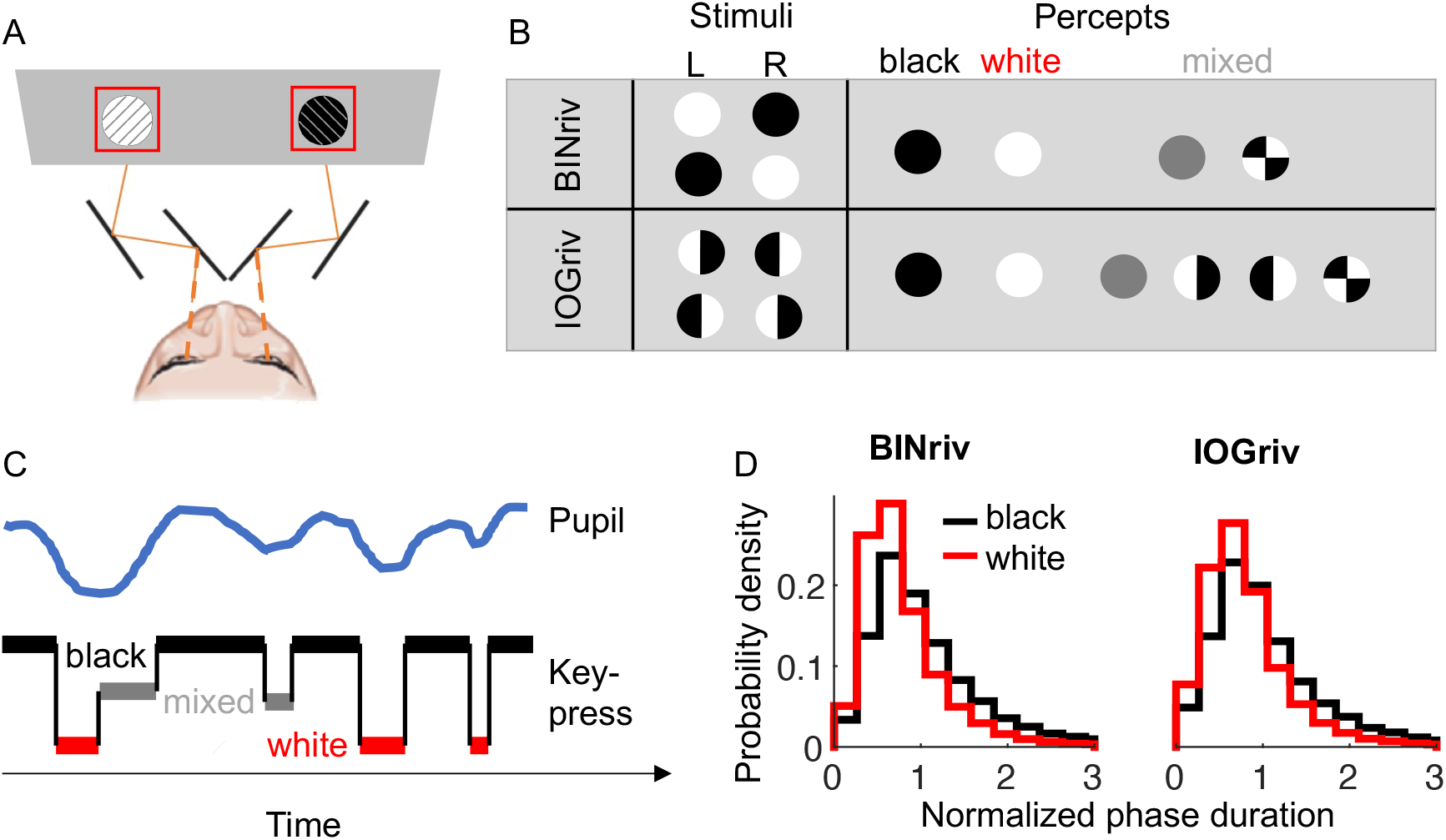
Dichoptic stimulation and rivalry dynamics. A: Schematics of the stimuli (white or black patches overlayed with orthogonal thin lines), presented dichoptically through a four-mirror stereoscope. B: Possible stimulus configurations (thin lines omitted) and perceptual outcomes for binocular rivalry (BINriv) and interocular grouping rivalry (IOGriv). C: Example traces from a segment of the experiment, where participants used keypresses to report the dominant percept (square wave) and we recorded pupil size oscillations (green wave). D: Probability density function of the normalized phase durations for exclusive dominance of white or black disk percepts in binocular rivalry and interocular grouping rivalry.

For interocular grouping rivalry, the same stimuli (white/black disks with thin lines) were split vertically and the two halves shown to the two eyes (Figure 1B). Possible percepts were: exclusive dominance of the white or the black disk (grouped interocularly), monocular percepts (half white half black disks, as shown to the left or to the right eye) and fusion or piecemeal percepts.

Stimuli were presented continuously for 3 min long trials. Trials were separated by 60 s long pauses with only the fixation point shown against the background. During this time, participants reported their perception of afterimages; analyses of this behavior will be reported in a separate publication. On each trial, a different combination of disk color and line orientation was presented to each eye; combinations were varied pseudo-randomly across trials. Participants continuously reported perception by keeping one of three keys pressed: right or left arrows to report dominance of the stimulus with clockwise or counterclockwise tilted lines, or the down-arrow key any time dominance of either stimulus was incomplete (i.e. monocular percepts in interocular grouping rivalry, piecemeal and fusion events were not distinguished in our paradigm).

Dominance phase distributions were adequately captured by a typical gamma distribution (Levelt, 1967), with shape *α* and scale *β* parameters for binocular rivalry: *α* = 2.62; *β* = 0.32 and for interocular grouping rivalry: *α* = 2.50 and *β* = 0.34 in Equation 1. The goodness-of-fit (coefficient of determination R^2^) was 0.94 for binocular rivalry and 0.97 for interocular grouping.

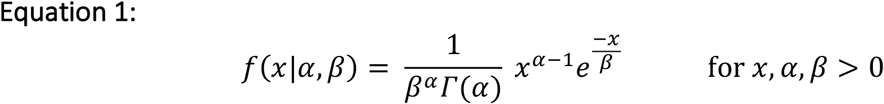

Where *Γ* is the gamma function and *x* the number of dominance phases.

Binocular rivalry and interocular grouping rivalry were both tested in three conditions: no attentional cue, black percept cued or white percept cued. In the latter, a “D” or a “B” letter was displayed at the beginning of the trial, cueing participants to endogenously focus their attention on the black or white disk throughout the rivalrous alternations.

We applied a fully randomized factorial design, where each subject completed 32 trials, divided into 8 sessions of four trials each, one for each combination of rivalry type and attentional cueing, distributed over two days.

We also set-up a simulated rivalry stimulus, where a single white, black or half white and half black disk (the latter simulating mixed percepts) was shown monocularly to one eye (right or left in separate trials), alternating with phases of 2.5 ± 0.01 s. Four trials of this stimulus were always run at the beginning of the experiment, to help participants familiarize with the task.

### Eye tracking data acquisition and analysis

During rivalry and simulated rivalry, we monitored pupil diameter and two-dimensional eye position with an infrared camera (EyeLink 1000 system, SR Research, Canada) mounted below the monitor screen and behind the stereoscope. EyeLink data were streamed to the main computer through the EyeLink toolbox for Matlab (Cornelissen et al., 2002) and thereby synchronized with participant’s keypresses. Pupil diameter measurements were transformed from pixels to millimeters using an artificial 4 mm pupil positioned at the approximate location of the subject’s eye.

Pupil and gaze tracking data consisted of 180 × 1000 (180 seconds at 1000 Hz) time points. These included signal losses, eyeblinks and other artifacts, which we cleaned out by means of the following steps (all implemented with in-house Matlab software):

- Identification and removal of gross artifacts: removal of time-points with unrealistically small or large pupil size (more than 1.5 mm from the median of the trial or <0.2 mm, corresponding to blinks or other signal losses).
- Identification and removal of finer artifacts: identification of samples where pupil size varied at unrealistically high speeds (>0.05 mm per second, beyond the physiological range) and removal of the 80 ms epoch surrounding this disturbance.
- Removal of any linear trend by fitting a linear function to pupil data from each 180 s-long trial.

After this cleaning procedure was applied, we verified fixation stability by measuring the dispersion of eye position samples around the mean of each trial as the bivariate confidence ellipse area (BCEA), defined as:

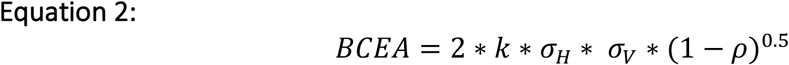

Where *k* is the confidence limit for the ellipse, *σH* and *σV* are the standard deviation of eye positions in the horizontal and vertical meridian respectively, ρ is the product-moment correlation of these two position components and *k*=1.14, implying that the ellipse included 68% (1-e^(−*k*)^) of the distribution. To test for possible differences in eye-movement patterns, we averaged the BCEA values across trials and entered these values in a 2×2 repeated measure ANOVA with factors: cueing condition (no cue/cueing) and rivalry type (binocular/ interocular grouping rivalry). No main effect or interaction was significant, suggesting that fixation was equally stable across all conditions and rivalry types (main effect of cueing condition: F_(1,37)_ = .22, p = .64, logBF = −.70; main effect of stimulus type: F_(1,37)_ = .16, p = .69, logBF = −.75; cueing condition by stimulus type interaction: F_(1,37)_ = .36, p = .55, logBF = −.50).

After cleaning, pupil data and continuous recordings of perceptual reports were down-sampled to 100 Hz, by taking the median of the retained time-points in non-overlapping time windows. If no retained sample was present in a window, that window was set to “NaN” (MATLAB code for “not a number”). Down-sampled pupil traces were finally parsed into epochs locked to each perceptual switch (when the subject changed perceptual report) and labelled according to the color of the dominant stimulus after the switch. Pupil time courses were averaged across epochs for each participant; further averaging across participants yields traces in Figure 2. In order to minimize the impact of pupil size changes unrelated to the perceptual switches, we also analyzed data after subtracting a baseline from each epoch, measured in the −1:−0.5 s interval preceding the switch. In order to compare pupil size across dominance phases, stimuli and attention conditions, we extracted an index of pupil size in each epoch by averaging its value in the −0.5:1 s interval around the switch. Note that shifting the intervals for pupil baseline, or skipping the baseline correction step, affected the size of pupil modulations but it did not change our conclusion on the effects of attention and stimulus type.

**Figure 2.**
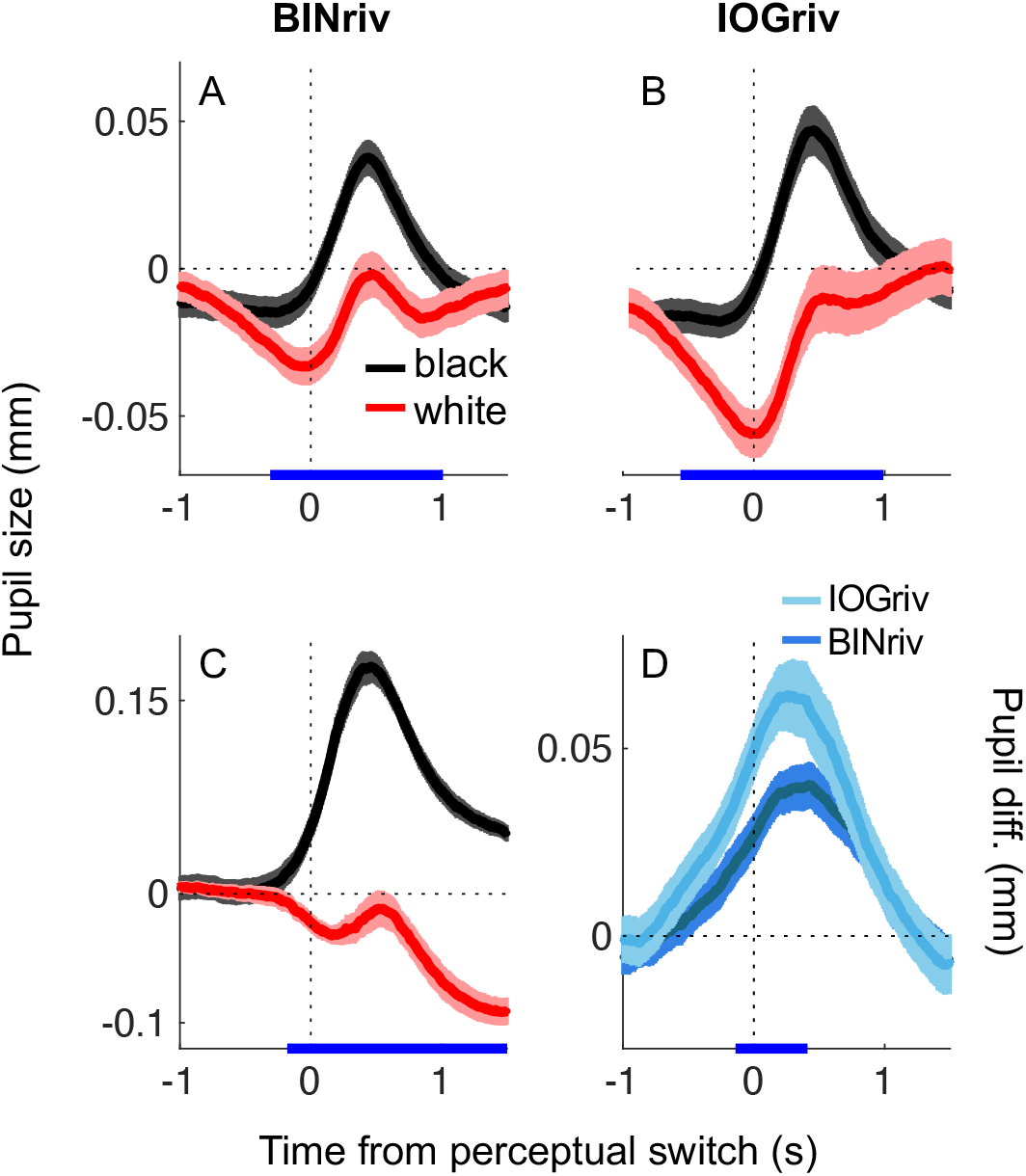
Pupil oscillations track perceptual alternations. A–C: pupil size traces aligned to perceptual switches towards exclusive dominance of a white disk or a black disk percept and averaged across phases, separately for binocular rivalry, interocular grouping rivalry and a simulated rivalry (physical alternation of the two disks in monocular viewing). D: average difference between pupil traces for white or black dominance phases, computed in individual participants and then averaged. In all panels: shadings report s.e. across participants and the blue marks on the x-axis highlight timepoints where pairwise comparisons between traces are significant (p < 0.05 FDR corrected).

In order to quantify the effect of attention on both perceptual reports and pupil measurements, we computed indices of attentional modulation (AMI) for comparing perceptual and pupil measures in cueing versus in no-cueing trials. Specifically, we used Eq. 3–4, where PROP is the total dominance time of the white and black disk percepts divided by total testing time and PUPDIFF is the average pupil size difference between black and white disk dominance phases.

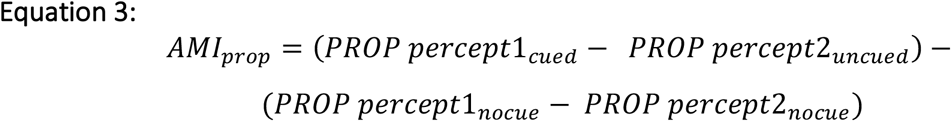

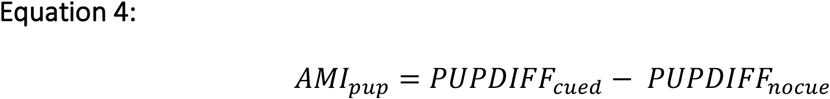

Modulation indices for the contrast manipulations in the control experiment were computed using the same equations and replacing “cued” with “contrast enhanced”. For the sake of clarity, we chose to quantify perceptual reports using dominance proportions; however, the same conclusions could be drawn analyzing mean phase durations instead.

### Statistical approach

Significance was evaluated using both p-values and log-transformed JZS Bayes Factors computed with the default scale factor of 0.707 (Wagenmakers et al., 2012). The Bayes Factor is the ratio of the likelihood of the two models H1/H0, where H1 is the experimental hypothesis (effect present) and H0 is the null hypothesis (effect absent). By convention, a base 10 logarithm of the Bayes Factor (logBF) > 0.5 is considered substantial evidence in favor of H1, and logBF < −0.5 substantial evidence in favor of H0. Bayesian ANOVAs were run in JASP and the corresponding Bayes Factors represent the change from prior to posterior inclusion odds (BFinclusion) computed across matched models. We estimated the internal consistency of our parameter estimates by split-half reliability. Each parameter was estimated twice per participant, on half the dataset (odd and even trials) and we evaluated the correlation of the two sets across participants. Finally, we evaluated the significance of behavioral attentional effects at the single subject level with a bootstrapping approach, by resampling (10,000 times, with reinsertion) dominance phases in cueing and no-cueing conditions, applying equation 3, computing the proportion of samples where the attentional modulation index was larger than 0 or smaller than 0 and assigning the significance for p < 0.025.

### Control experiment

A control experiment was performed after the conclusion of the study, with the aim of estimating the sensitivity of pupil size measurements to manipulations of stimulus strength. The original set-up was unaccessible at the time of testing and we replicated the conditions of the main experiment as closely as possible in another set-up, using the same eyetracker (EyeLink 1000 system, SR Research, Canada), similar mirror stereoscope and a computer that ensured equal performance. Specifically, stimuli were generated with the PsychoPhysics Toolbox routines (Brainard, 1997) or MATLAB (MATLAB r2010a, The Math Works) housed in a Mac Pro 4.1, and displayed on a 52.5 cm-wide LCD screen, with 12.3 cd/m^2^ gray background as in the main experiment. We tested binocular rivalry in six conditions: the no-cue and white-cued conditions were the same as in the main experiment; in addition, we tested four conditions where the Michelson contrast of the white disk stimulus was increased by 25%, 50%, 100%, 200% (luminance respective values: 35 cd/m^2^,43 cd/m^2^,77 cd/m^2^, 150 cd/m^2^). Each condition was tested in four trials, and all data were collected over a single session.

## Results

We analysed perceptual alternations and pupil oscillations during binocular and interocular grouping rivalry in two conditions, with and without attentional cueing.

In no cueing conditions, pupil diameter reliably tracked perceptual alternations between a white and a black disk presented dichoptically, either one disk per eye generating binocular rivalry, or each disk split vertically between eyes generating interocular grouping rivalry (Figure 1). In spite of constant stimulation (hence constant luminance), pupils were relatively dilated when participants reported seeing black, compared to when they reported seeing white (Figure 2A-B, black vs red line), in both types of rivalry. Table 1 (rightmost column) show the statistics computed for average pupil size measurements, highlighting a strong effect of the factor “dominant percept” (white/black) and no effect of rivalry type (binocular/interocular grouping rivalry) or interaction between the two. For comparison, we acquired pupil data from a simulated rivalry condition where the white and black disk were physically alternated in monocular viewing (Figure 2C). Pupil oscillations during binocular and interocular grouping rivalry were a sizable portion of those during these physical luminance alternations, averaging respectively 28.8 ± 6.58% (mean ± S.E.M.) and 31.3 ± 7.86% of the pupil oscillations observed during simulated rivalry. Note the tendency for oscillations in interocular grouping rivalry to be more pronounced than for binocular rivalry (Figure 2D), which might reflect small differences in perceptual reporting strategies. Given that exclusive white/black dominance are much rarer events in interocular grouping than binocular rivalry (see below) participants may have spontaneously adopted a stricter criterion for reporting these percepts, resulting in pupil modulations being less contaminated by unreported partial dominance events. Figure 2D also highlights that pupil size modulations consistently started before the perceptual switch, and this was more pronounced in the rivalry conditions than in the simulated rivalry (significant pupil difference started 310 ms before the switch in binocular and 560 in interocular grouping rivalry, compared with the 179 ms latency for simulated rivalry). This may reflect the graded nature of rivalry transitions, which may delay change detection in rivalry compared with the sharp transitions used in the simulated rivalry condition.

**Table 1.**
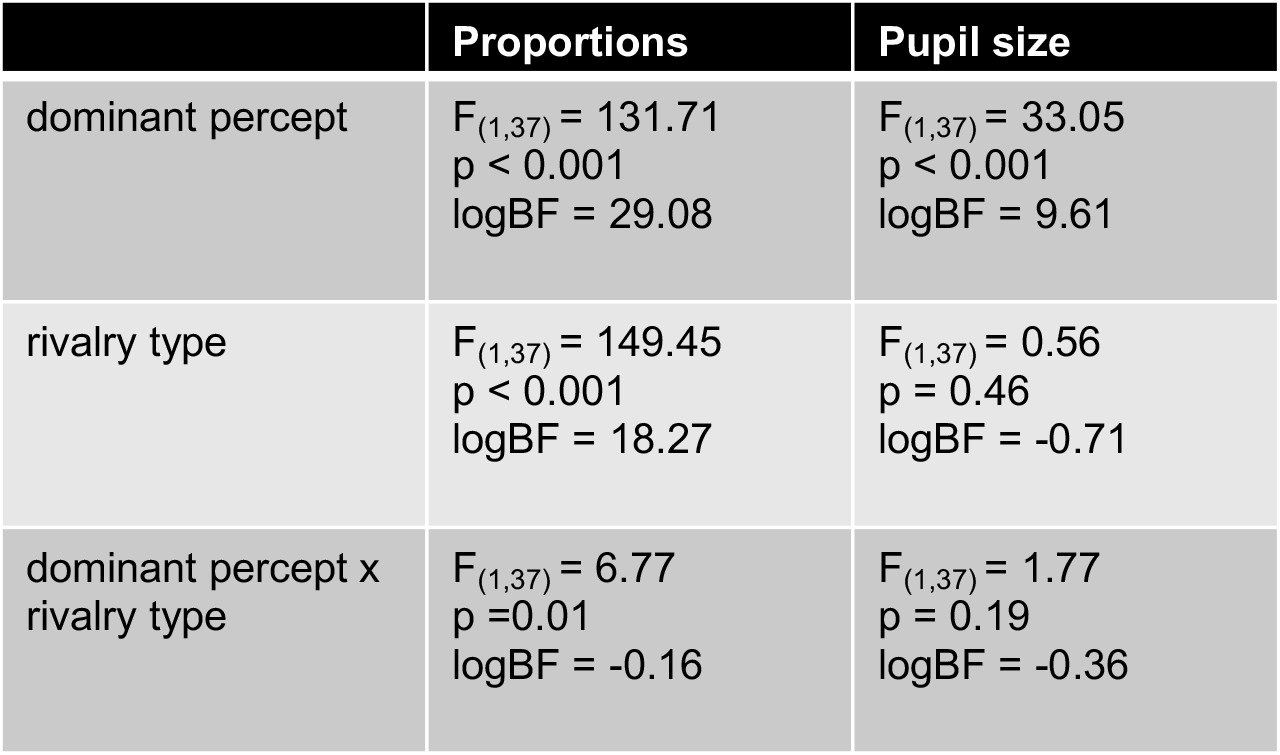
Two-way ANOVA for no-cue results, with factors: percept type (white/dark disk) and rivalry type (binocular/interocular grouping rivalry).

The analysis of behavioural reports in the no cueing condition (dashed lines in Figure 3) showed a net predominance of black disk percepts with respect to the white ones (main effect of dominant percept in Table 1), marginally more pronounced for interocular grouping rivalry. In line with the modified Levelt’s propositions and results by Qiu et al., (2020), this can be explained by the higher Michaelson contrast of the black disk stimulus (recall that stimulus luminance was set to the limits of the screen to evoke the largest possible pupil modulations).

**Figure 3.**
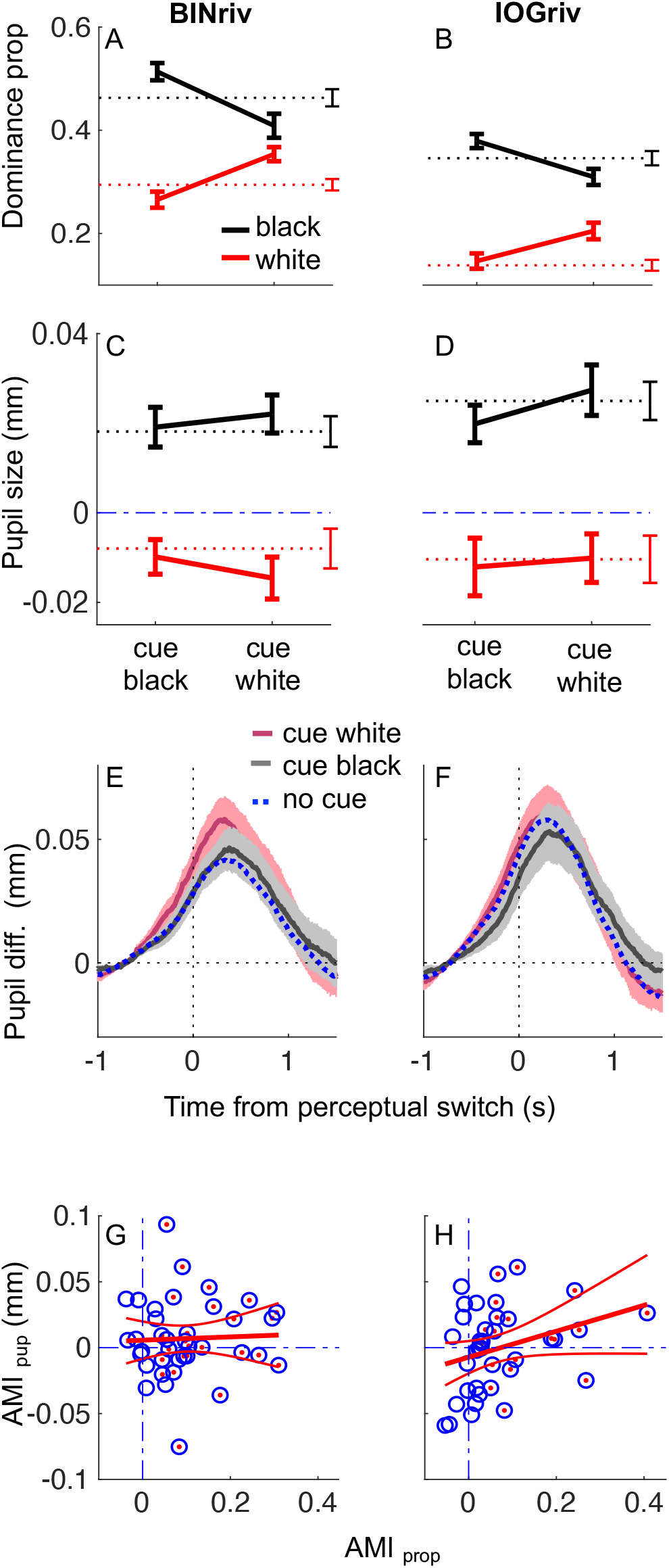
Attention cueing affect perceptual alternations but not pupil oscillations. A-D: perceptual dominance (A-B) and baseline corrected average pupil size (C-D) for exclusive white or black disk percepts, without attentional cueing (dashed lines) or when the white or the black disk percept were cued (continuous lines). Error bars report s.e. across participants. E-F: timecourse of the difference between pupil traces for white or black dominance phases, computed in individual participants and then averaged for each cueing condition. Shaded areas show s.e. across participants. G-H: individual participants’ attentional modulation indices for perceptual dominance (x-axis) and pupil size (y-axis), computed with Equations 3–4 respectively. Dash-dot blue lines mark the x=0 and y=0 lines, indicating no effect of attention cueing. Each circle reports results from one participant; red dots highlight participants with a significant attentional modulation index for perceptual dominance. Red lines show the best fitting line and its 95% confidence intervals. In all panels, the left column reports results for binocular rivalry and the right for interocular grouping rivalry.

However, dominance of the black percept cannot logically explain the pupil modulations; moreover, while both black dominance and pupil size modulations varied across participants, the two were reliably uncorrelated (binocular rivalry: r = −.15, p = .37, logBF = −.72, interocular grouping rivalry: r = .10, p = .55, logBF = −.82).

Figure 3A-B also show that exclusive percepts were much rarer in interocular grouping rivalry compared to binocular rivalry (main effect of rivalry type on dominance proportions in Table 1), reflecting the response mapping we used; for interocular grouping rivalry, mixed reports included epochs where the individual monocular images in the left and right eye dominated.

Having established pupil size as a marker of perceptual dominance in both binocular and interocular grouping rivalry, we proceeded to assess the impact of attention cueing on perceptual alternations and pupil size oscillations (continuous lines in Figure 3).

As expected, perceptual dominance of the cued stimulus was enhanced, resulting in a significant interaction between dominant percept (white/black) and cued percept (cueing white/cueing black, Table 2). We summarized the effect of attention with an attentional modulation index (Eq. 3 in Methods), which was in the order of 10% for both types of rivalry (single subject data are shown on the abscissas of Figure 3G-H and Figure 4A shows averages). The effect was statistically reliable at the group level (Table 3) and, in most cases, at the individual subject level (bootstrapped attentional modulation indices were significantly higher than zero in 27/38 or 20/38 participants for binocular and interocular grouping rivalry respectively, highlighted with a red dot in Figure 3G-H; it was significantly lower than zero in only 2/38 participants for interocular grouping rivalry and in no participant for binocular rivalry). Attentional modulation indices were correlated between binocular rivalry and interocular grouping (r = .61, p < .001, logBF = 2.77), suggesting that they measure a relatively stable feature of our participants. In line with this, we found no indication that interocular grouping rivalry was more affected by attention than binocular rivalry; if anything, there was a small tendency for an opposite effect (Table 3, coherent with the significant three-way interaction between dominant percept, cued percept and rivalry type for dominance proportions reported in Table 2).

**Table 2.** Three-way ANOVA for attention cueing results, with factors: percept type (white/dark disk), cueing (white/dark cued), rivalry type (binocular/interocular grouping rivalry).

|  | Proportions | Pupil size |
| --- | --- | --- |
| dominant percept | $F_{(1,37)} = 155.50$<br>$p < 0.001$<br>$\log BF = 32.28$ | $F_{(1,37)} = 41.48$<br>$p < 0.001$<br>$\log BF = 20.02$ |
| rivalry type | $F_{(1,37)} = 82.96$<br>$p < 0.001$<br>$\log BF = 25.55$ | $F_{(1,37)} = 0.45$<br>$p = 0.50$<br>$\log BF = -0.82$ |
| cued percept | $F_{(1,37)} = 5.56$<br>$p = 0.02$<br>$\log BF = -0.79$ | $F_{(1,37)} = 1.33$<br>$p = 0.26$<br>$\log BF = -0.82$ |
| dominant percept<br>x rivalry type | $F_{(1,37)} = 0.88$<br>$p = 0.35$<br>$\log BF = -0.61$ | $F_{(1,37)} = 0.08$<br>$p = 0.77$<br>$\log BF = -0.66$ |
| dominant percept<br>x cued percept | $F_{(1,37)} = 32.11$<br>$p < 0.001$<br>$\log BF = 11.26$ | $F_{(1,37)} = 1.73$<br>$p = 0.20$<br>$\log BF = -0.55$ |
| rivalry type<br>x cued percept | $F_{(1,37)} = 0.27$<br>$p = 0.63$<br>$\log BF = -0.68$ | $F_{(1,37)} = 1.72$<br>$p = 0.20$<br>$\log BF = -0.61$ |
| dominant percept<br>x rivalry type<br>x cued percept | $F_{(1,37)} = 5.59$<br>$p = 0.02$<br>$\log BF = -0.25$ | $F_{(1,37)} = 0.04$<br>$p = 0.85$<br>$\log BF = -0.71$ |

**Figure 4.**
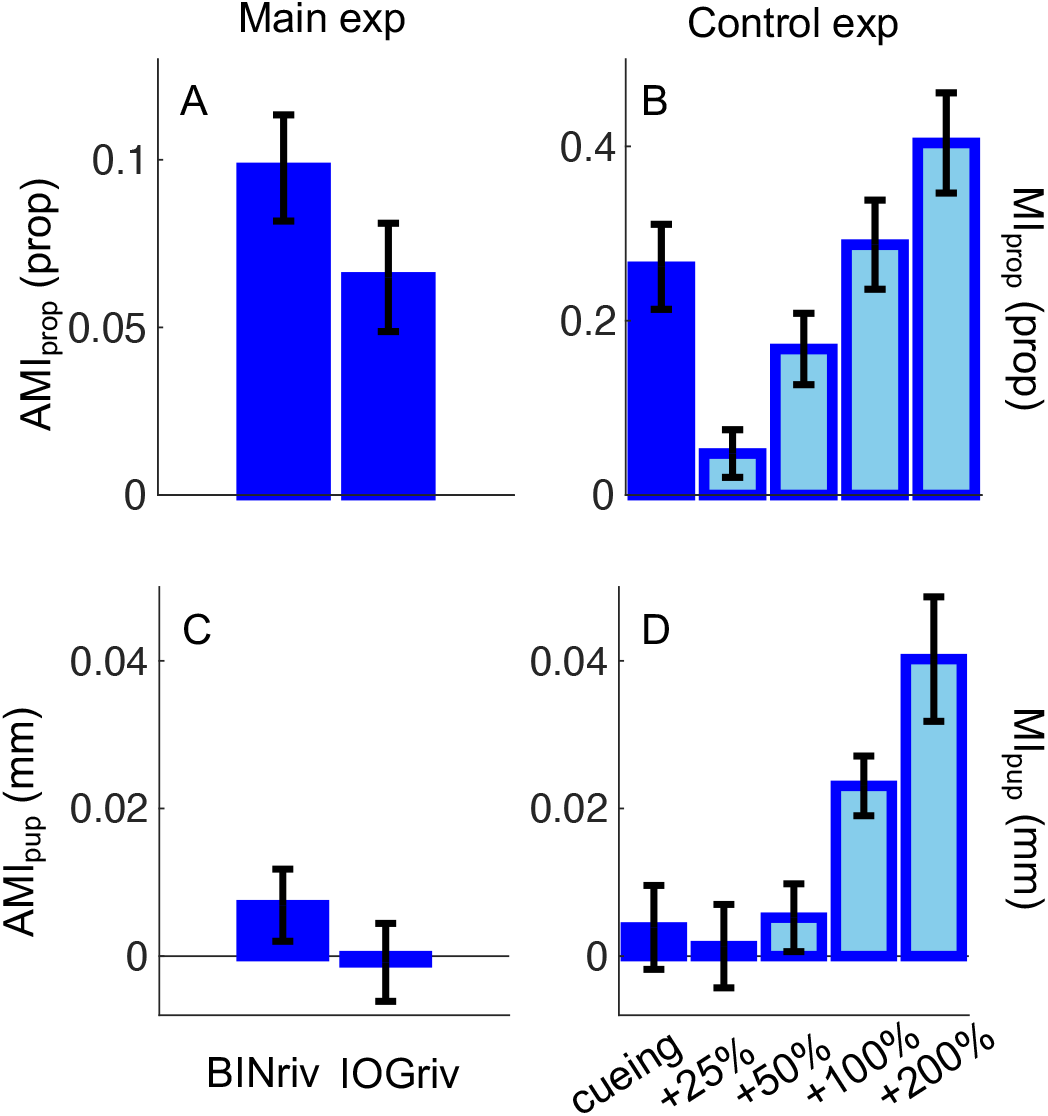
Effects of attention cueing vs. enhancing contrast. Modulation indices for perceptual dominance (A-B) and baseline corrected pupil size (C-D) computed by contrasting cued vs. uncued percepts in the main experiment (left column) and cueing or enhancing contrast of the white stimulus (right column). In all panels, error bars report s.e. across participants.

**Table 3.** T-tests on attentional modulation indices

|  | AMI Proportions | AMI Pupil size |
| --- | --- | --- |
| binocular rivalry | $t_{(37)} = 6.20$<br>$p < 0.001$<br>$\log BF = 4.67$ | $t_{(37)} = 0.16$<br>$p = 0.88$<br>$\log BF = -0.75$ |
| interocular grouping | $t_{(37)} = 4.01$<br>$p < 0.001$<br>$\log BF = 1.99$ | $t_{(37)} = 1.22$<br>$p = 0.23$<br>$\log BF = -0.45$ |
| binocular rivalry<br>vs.<br>interocular grouping | $t_{(37)} = 2.36$<br>$p = 0.02$<br>$\log BF = 0.31$ | $t_{(37)} = 0.78$<br>$p = 0.44$<br>$\log BF = -0.63$ |

Based on the assumption that attention cueing boosts perceptual dominance by enhancing the effective strength of the cued percept, we expected to find an enhancement of the pupil oscillations accompanying perceptual alternations. For example, we predicted that dilations concurrent with black percept dominance would be increased when cueing black. This should have resulted in larger differences in pupil size during black vs. white percepts under attentional cueing as compared to no cueing conditions. However, contrary to this prediction, we found that the magnitude of the pupil oscillations was unaffected by attentional cueing. Figure 3C-D plots the mean pupil size over a fixed interval [−0.5:1s] around the perceptual switch for each percept type and attention cueing condition; the format is the same as for Figure 3A-B, but clearly there are no systematic differences when the white or the black percept were cued or compared to the no cueing condition (dashed lines). Figure 3E-F shows the pupil difference traces, which were comparable across attentional conditions, implying that pupil oscillations reliably tracked perceptual alternations in all conditions, but did not change when one percept was cued. As shown in Table 2 (rightmost column), both the main effect of cued percept and the interaction between dominant percept and cued percept are non-significant, with log Bayes Factors < −0.5 (Table 2). Coherently, the attentional modulation indices (computed with Equation 4) were distributed around zero for all participants (as shown on the ordinates of Figure 3G-H, Figure 4C and Table 3). Even selecting the subsample of participants who showed a significant behavioural effect of attention cueing (marked by red dots in Figure 3G-H), the pupil attentional modulation remained non-significantly different from zero (t(26) = .84, p = .40, logBF = −.56 for binocular rivalry and t(19) = .75, p = .46, logBF = −.53 for interocular grouping rivalry).

Thus, our results show disagreement between pupillometric and behavioural measures of perceptual dominance. Only perceptual alternations were affected by attention cueing, not the accompanying pupil oscillations. Could this be due to lack of sensitivity of pupillometry? We gathered several pieces of evidence against this possibility.

First, Bayesian statistics (log Bayes Factors associated with effects of cueing were below −0.5, Tables 2 and 3) indicate that we do not just lack evidence in support of an effect of attention, we actually have robust evidence in support of the opposite: that attention cueing does not affect pupil size.

Second, the reliability of pupil measurements was high, approaching that of the behavioural measurements (test-retest reliability for binocular rivalry: dominance proportions 0.84, logBF = 8.51 and pupil size difference: 0.79, logBF = 6.71; interocular grouping rivalry: dominance proportions 0.89, logBF = 10.81 and pupil size difference 0.84, logBF= 8.46).

Third, pupil measurements were sensitive enough to report the slight unbalances between eyes observed in our set of (non-amblyopic) participants. This was shown by splitting the same set of perceptual phases (binocular rivalry with attention cueing) in two ways: according to whether the reported percept was cued or un-cued, and according to whether it matched the stimulus presented in the dominant or non-dominant eye. This measure confirmed that pupil size was insensitive to attention cueing (pupil modulations were not different when the cued or un-cued stimulus was perceived, t_(37)_ = .18, p = .86, logBF = −.75) but it did report eye-dominance (pupil modulations being larger in phases where percepts matched the stimulus in the dominant eye, t_(37)_ = 2.15, p = .04, logBF = .14).

Fourth, and finally, the results of a control experiment showed that pupil size modulations reliably track changes in stimulus strength of a size compatible with those simulating the effects of attention cueing on perceptual dominance (Figure 4, right column). This was tested in a separate cohort of participants, where we repeated the attentional manipulation (comparing trials where the white disk was cued with no-cue trials) and, in separate no-cue trials, we manipulated the physical strength of the white disk. Table 4 shows paired t-tests comparing the attentional modulation index for white-cueing and the modulation index for each of the contrast enhancement conditions. The effect of cueing the white disk could be effectively simulated (in no-cue conditions) by doubling the contrast of the white disk, i.e. a 100% increase. However, the pupil behaved very differently in the two cases. Doubling the contrast of the white disk was sufficient to reliably increase of the pupil oscillations, but pupil oscillations were not enhanced when the white percept was cued compared to the no-cue case, replicating the results of our main experiment.

**Table 4.**
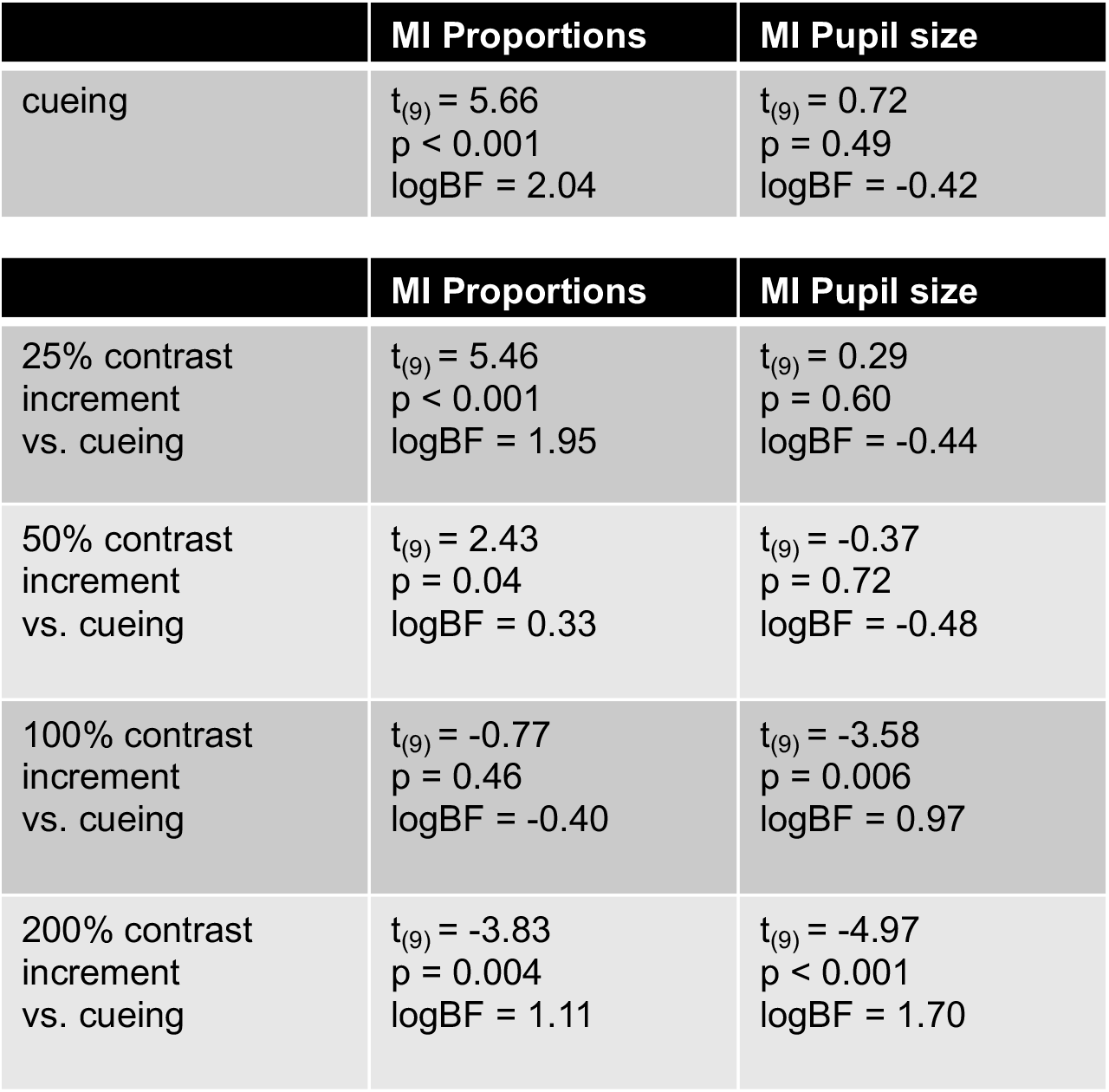
Modulation indices from the control experiment: effect of attention cueing and comparison of attention cueing vs. contrast enhancement by 25, 50, 100 and 200%

### Analysis of mixed percepts

Figure 5 reanalyses the results from the main experiment focusing on reports of mixed percept (defined as anything but the exclusive dominance of a white or a dark disk percept). We found that attention cueing did not affect the proportion of mixed percepts in binocular rivalry, which averaged .24±.02 and .23±.02 for no cueing and cueing conditions (t_(37)_ = .86, p = .40, logBF = −.61). In interocular grouping rivalry, the difference was also non-significant but there was a trend towards reduced mixed reports in the cueing condition (.52±.02 and .48±.02 for no cueing and cueing respectively, t_(37)_ = 1.78, p = .08, logBF = −.14). Based on this observation, we cannot exclude the possibility that attention cueing may have promoted interocular grouping. Figure 5 C-D show pupil traces aligned to the onset of mixed percepts, separately for no cueing and the two attention cueing conditions; note that this is conceptually equivalent to analysing data for exclusive dominance phases aligned to their offset, rather than the onset (same conventions as in Figure 3 E-F). A difference between cueing conditions is apparent for binocular rivalry, suggesting enhanced pupil dilation when the white percept was cued. No such effect is observed for interocular grouping rivalry, possibly due to the heterogeneous nature of mixed percepts.

**Figure 5.**
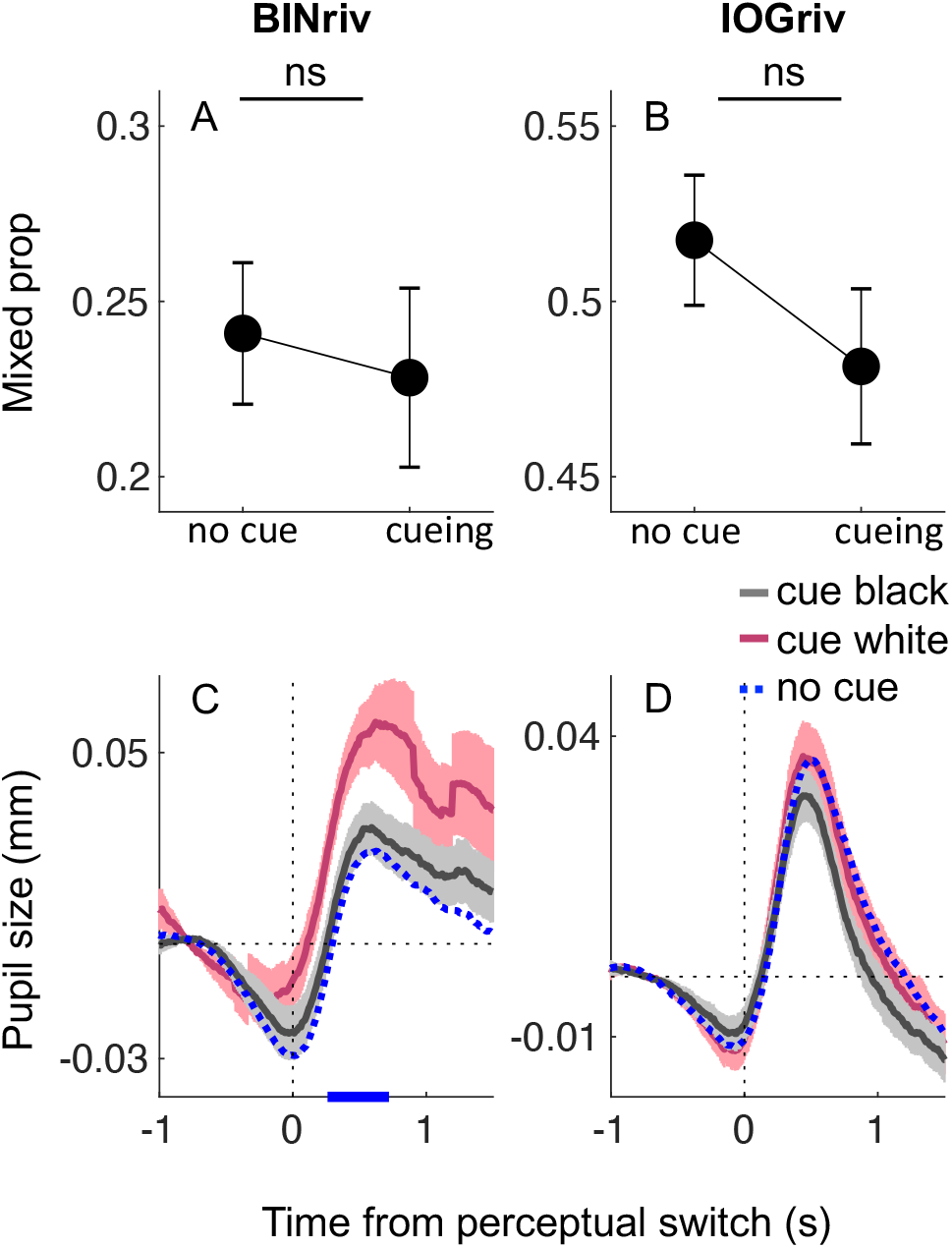
Effects of attention cueing on mixed percepts. A: proportion of mixed percepts in no cueing or cueing conditions (collapsed across white and black cued). Error bars report s.e. across participants B: pupil traces during mixed percepts in the three cueing conditions. Shaded areas show s.e. across participants and blue marks on the x-axis highlight timepoints where pairwise comparisons between the white and black cueing conditions are significant (p < 0.05 FDR corrected).

This difference between the two rivalry types may suggest that the cueing effect is specifically related to fusion percepts (grey disk percepts), which likely represented a minor percentage of mixed reports in interocular grouping rivalry (the largest majority being monocular half white, half black disk percepts). The relative pupil dilation may indicate that fusion events were perceived as grey disks of a darker shade during white cueing than during dark cueing or no cueing. There are at least two reasons why this could happen. One possibility is that white cueing indeed enhanced the effective strength of the white stimulus, implying that the black stimulus strength had to reach a higher threshold before a white-dominance report switched to a mixed report. If this is the case, however, it is unclear why such difference in effective strength should not show in pupil traces during phases of exclusive white dominance (Figures 3 and 4). Another possibility is that cueing may have affected decision criteria so that fusion percepts reported as mixed were generally darker under white cueing (effectively prolonging exclusive white-dominance phases) than under black cueing conditions.

## Discussion

We used pupillometry to investigate the effects of endogenous attention on binocular rivalry and interocular grouping rivalry.

We confirmed that pupil size tracks perceptual oscillations during binocular rivalry, in spite of constant luminance stimulation, and we extended this observation to interocular grouping rivalry. This is consistent with the large body of work suggesting that the subcortical circuit generating the pupillary light response can be modulated by perceptual signals (Binda and Murray, 2015; Binda and Gamlin, 2017; Mathot, 2018). Our finding that similar pupillary modulations accompany interocular grouping rivalry constrains the origin of the modulatory signals to visual cortical areas (they must hold a representation of stimulus brightness) with access to binocular information (they must be able to combine information from the two eyes). We found that manipulating endogenous attention reliably affected perceptual alternations, enhancing dominance of the cued percept during binocular rivalry, in line with previous work (Meng and Tong, 2004; Mitchell et al., 2004; Chong et al., 2005; Hancock and Andrews, 2007; Paffen and Alais, 2011). To our knowledge, this is the first report of the effects of manipulating attention in interocular grouping rivalry. We found that attention cueing had the same or slightly smaller effects on interocular grouping rivalry as on binocular rivalry. This suggests that eye-based and pattern-based competition are similarly permeable to endogenous attention; it also suggests that different degrees of attentional control (as observed, for example, comparing Necker cube vs. binocular rivalry, Meng and Tong, 2004) may be related to differences in stimulus complexity rather than to the involvement of earlier (monocular) cortical representations.

Many have suggested that attention cueing acts by enhancing the perceptual strength of the cued signals (Carrasco, 2011). This is in line with evidence that focusing attention at a spatial location or feature enhances its representation in early visual cortex, as measured with EEG (Hillyard and Anllo-Vento, 1998; Di Russo et al., 2003; Wang et al., 2007; Kelly et al., 2008; Khoe et al., 2008; Mishra and Hillyard, 2009), fMRI (Saenz et al., 2002; Liu et al., 2005; Boynton, 2009; Pestilli et al., 2011) or indexed by enhanced pupillary response to light stimuli at the attended location (Binda and Murray, 2015). Transferring this knowledge to the context of rivalry, we expected that cueing attention to one of the rivaling percepts would enhance its effective strength and thereby increase its dominance. Using pupillometry, we intended to indirectly index this phenomenon. We established that the magnitude of pupil oscillation accompanying rivalry is sensitive to effective stimulus strength as set by ocular dominance (control analysis of binocular rivalry data from the main experiment) or physical contrast changes (control experiment). On this basis, we predicted that attention cueing would have a similar effect as physical contrast enhancement, namely an amplification of pupil oscillations. However, we obtained evidence against this prediction, as pupil oscillations during periods of exclusive dominance were reliably unaffected by attention cueing.

The simplest way to explain this negative finding is putting it down to insufficient sensitivity of the pupillometric measurements. However, our reliability analysis, Bayesian statistics and results from a control analysis and a control experiment all coherently speak against this possibility. We therefore speculate on a few logical alternatives.

During binocular rivalry, most of the time is spent in exclusive dominance, where competition between rivaling stimuli is resolved, leaving only one visible stimulus and no distracter. In these conditions, attention may be automatically driven to the dominant stimulus (Li et al., 2017), leaving little space for endogenous re-directing of attention. Although this is consistent with attention affecting early visual processing in markedly different ways at the onset of rivalry vs. for non-rivaling stimuli (Khoe et al., 2008; Mishra and Hillyard, 2009), the model by Li et al. (2017) does not explicitly account for the small but reliable effects of attention cueing on perceptual alternations during rivalry. To account for these, one possibility is assuming that attention cueing primarily affects rivalry when the competition between stimuli is unresolved, namely in the brief times marking transitions between exclusive dominance phases, when the depth of suppression decreases (Alais et al., 2010). This idea has been suggested previously and supported by the observation that exogenous cues are mostly effective when presented near the end of individual dominance periods (Dieter et al., 2015). In this scenario, we could reconcile our behavioral and pupillometry results assuming that attention enhances the strength of cued percepts only in short intervals near perceptual switches, not during the entire dominance phases. This could be consistent with our observation that the only effects of cueing over pupil traces could be observed in a brief interval during mixed percepts.

An alternative possibility is that attention cueing affects rivalry dynamics by acting on a stimulus representation that is not represented in pupil dynamics. Available evidence is consistent with pupil size integrating a cortical representation of stimulus brightness (e.g. one that oscillates, tracking rivalry dynamics), but we lack direct knowledge on the level at which such visual representation is generated and fed into the pupil control circuit (Binda and Gamlin, 2017). On the other hand, evidence indicates that rivalrous perception is orchestrated by the interplay of fronto-parietal and occipital regions, which participate in different degrees depending on details of the stimulus and task (Logothetis et al., 1996; Sterzer and Kleinschmidt, 2007; Sterzer et al., 2009). It is possible, then, that attention affects competition after the stage where visual representations are fed to pupil control – whether this needs to be a decisional stage or still a sensory representation cannot be determined based on the available research.

This is not the first case where we find that pupillary responses are independent of physical luminance and yet inconsistent with perceptual judgments (Benedetto and Binda, 2016; Turi et al., 2018; Pome et al., 2020; Tortelli et al., 2020, 2021). These inconsistencies were generally explained by calling decisional factors into the picture, as these may bias or add variability to perceptual reports while leaving pupil size unaffected (Tortelli et al., 2021). That contextual factors other than physical luminance affect pupil size and perception similarly but independently – if it proves recurrent and reliable across paradigms – might call for an updated model of pupil control. It might suggest that separate processing pathways support perception and pupil control, in analogy (or perhaps in overlap) with the separate pathways supporting vision for perception and vision for action (Goodale and Milner, 1992).

In conclusion, we find that pupil size oscillates in phase with perceptual oscillations during binocular and interocular grouping rivalry, implying cortical control. In spite of this, and in spite of reliable effects of attention cueing on behavioral reports, pupil size during periods of exclusive dominance does not show any modulation with attention. This introduces new constraints for models of attention in rivalry and pupil control: either attention cueing affects perception without enhancing the dominant percept, or we hold multiple representations of the dominant percept that independently regulate behavioral reports and pupil size and are differentially affected by attention cueing.

## Acknowledgments

This project has received funding from the European Research Council (ERC) under the European Union’s Horizon 2020 research and innovation programme (grant agreement No 801715 – PUPILTRAITS and No 948366 - HOPLA), from the Italian Ministry of Research and Universities under the funding schemes PRIN-2017 (grant MISMATCH) and FARE-2 (grant SMILY) and the French National Research Agency (ANR), AAPG 2019 JCJC (grant agreement ANR-19-CE28-0008, PlaStiC).

## Notes

### Competing Interest Statement

The authors have declared no competing interest.

